# Energetic implications of fMRI-based nodal complex network metrics: a complex picture unfolds across diverse brain states

**DOI:** 10.64898/2026.01.08.694967

**Authors:** Shirley Feng, Sean E. Coursey, Sara A. Nolin, Nicole R. Zürcher, Hsiao-Ying Wey, Jonathan R. Polimeni, Marjorie Villien, Anisha Bhanot, Bruce R. Rosen, Jacob M. Hooker, Jingyuan E. Chen

## Abstract

Functional MRI-based graph theory has provided profound insights into the brain’s functional organization, yet the neuroenergetic meaning of widely used graph-theoretical metrics remains poorly understood. Although resting-state research suggests a positive coupling between network topology and glucose metabolism, it remains unclear whether this relationship reflects a general principle of brain organization or a state-specific phenomenon. Here, we test the neuroenergetic interpretability of nodal graph-theoretical metrics by linking complex network topology to cerebral glucose consumption across diverse brain states. Leveraging simultaneous functional PET-MRI, we directly compare state-dependent fluctuations in glucose consumption and network topology during sensory, cognitive, and arousal conditions. We further assess metabolic-topological couplings in disease through a meta-analysis of resting-state FDG-PET and fMRI studies involving Alzheimer’s disease, Parkinson’s disease, major depressive disorder, and schizophrenia. Our results show that nodal graph-theoretical metrics exhibit state- and network-dependent metabolic associations, with coupling patterns diverging across experimental and disease contexts. Notably, frontoparietal and attentional networks show more conserved metabolic–topological coupling than other large-scale networks across states. These findings underscore a dynamic, complex interplay between metabolic demand and complex network organization, highlighting the need for a nuanced interpretation of the energetic underpinnings of nodal graph-theoretical metrics in health and disease.

## INTRODUCTION

Graph theory provides a powerful framework for characterizing the brain’s large-scale functional organization with functional magnetic resonance imaging (fMRI)^1–3^. In this framework, brain regions are nodes in the graph, and their functional connections are edges between nodes. This graph can be assessed with various complex network metrics, such as nodal hubness and network efficiency, that reflect distinct topological properties of neural systems^1,4,5^. Altered complex network metrics are consistently observed in neurological and psychiatric disorders^6–8^, where disrupted topology has been leveraged for disease identification with machine learning^9,10^. Task-based fMRI studies further show that network topology dynamically adapts to changing cognitive demands^11,12^. Despite their widespread use, the physiological processes underlying the topological patterns captured by graph theory remain poorly understood. Clarifying these processes is essential for anchoring such metrics in biophysiology and for guiding the interpretation of the extensive body of research built upon them.

Initial insights into the neuroenergetic basis of fMRI-derived graph-theoretical metrics have been achieved by linking them to regional glucose consumption, typically measured with ^18^F-Fluorodeoxyglucose (FDG) positron emission tomography (PET)^13–16^. These studies have primarily focused on nodal degree, a measure of how many connections a brain region has and thus its network centrality, testing whether regions with richer connectivity impose higher metabolic demands. Such investigations have consistently identified a positive association between nodal degree and glucose metabolism in the task-free resting-state brain, both across cortical regions^13–15^ and across individuals^16^.

Although these earlier findings suggest that highly connected regions may incur greater energetic costs, their generalizability beyond the resting state remains unclear. More critically, these associations are derived from static measures obtained at a single brain state, which may be partly driven by inherent inter-region or inter-subject variations. A more direct test of the neuroenergetic relevance of nodal graph-theoretical metrics may require examining whether they co-vary across states. Specifically, when a stimulus is applied or brain states shift, do changes in nodal network metrics mirror regional alterations in brain metabolism?

In this study, we directly compare how nodal graph-theoretical metrics and glucose metabolism fluctuate across a diverse range of brain states to determine whether these metrics carry consistent energetic implications. We leverage recent advances in FDG-based functional PET imaging (fPET-FDG), which enable the tracking of minute-scale glucose metabolism dynamics using a (bolus plus) constant infusion of radioactive tracers^17–20^. By combining fPET-FDG with simultaneous BOLD-fMRI on a hybrid PET-MRI scanner, we can measure dynamic changes in the functional cerebral metabolic rate of glucose (CMRglu) alongside fMRI-derived nodal graph-theoretical metrics within the same individuals and brain states^19^ (Fig. 1a). Using this multimodal paradigm, we examine four states: visual stimulation, finger tapping, working memory, and sleep to determine whether alterations in nodal network topology track changes in glucose metabolism.

**Figure 1.**
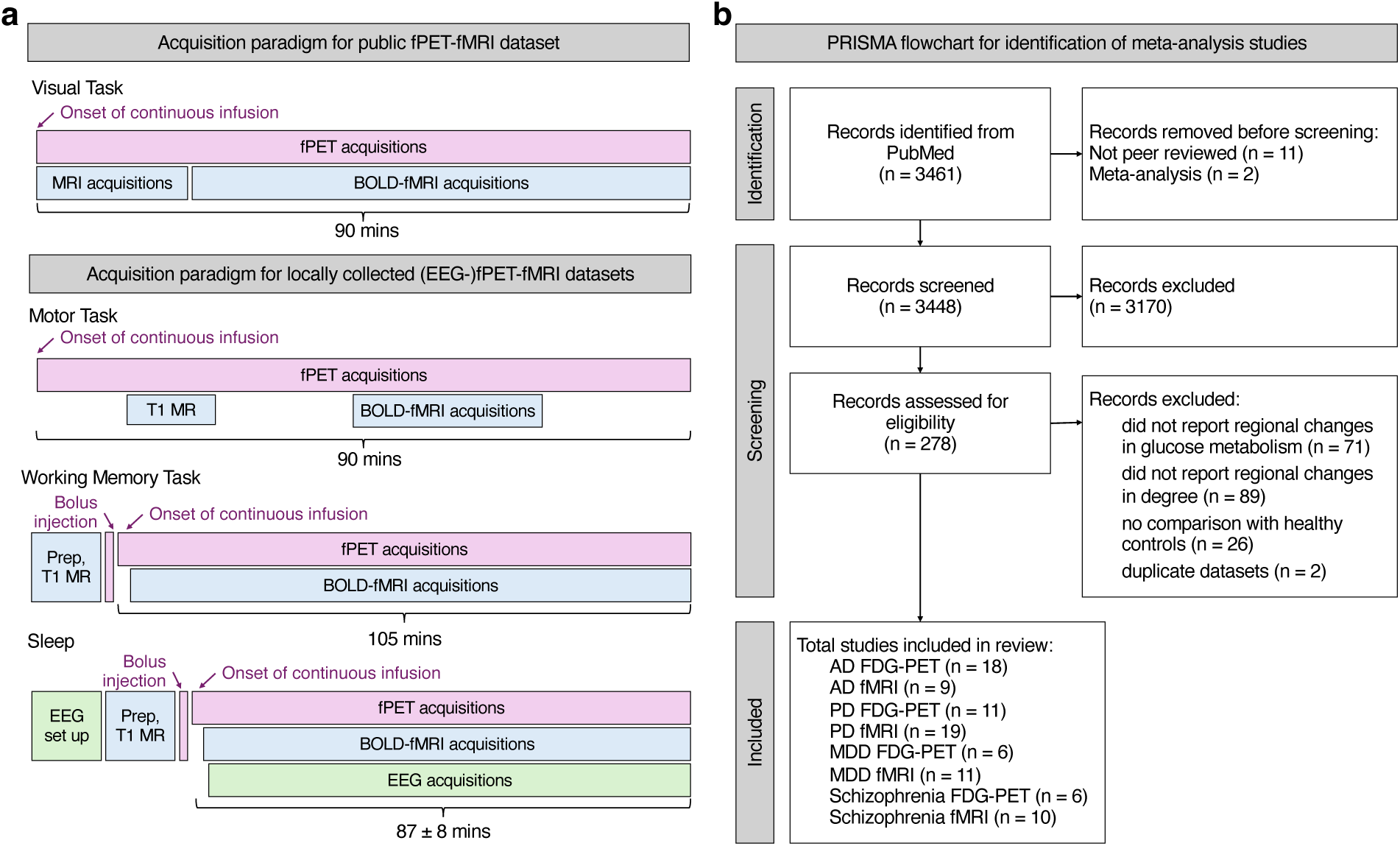
Summary of empirical fPET-fMRI and meta-analytic data used to probe state-related changes in nodal graph-theoretical metrics and glucose metabolism. (a) Experimental designs of simultaneous fPET-fMRI datasets examining sensory-, cognitive-, and arousal-related modulations of various nodal graph-theoretical metrics (BOLD-fMRI) and glucose metabolism (fPET-FDG). (b) PRISMA flowchart for the meta-analysis of disease-related alterations in nodal degree (ZOLD-fMRI) and glucose metabolism (static FDG-PET).

To extend these comparisons to clinical contexts, our empirical analyses are complemented by a meta-analysis of disease-related alterations in network topology and energetic demands (Fig. 1b). We synthesize reported changes in nodal degree (the most prevalent nodal metric in the literature) and glucose metabolism during the resting state in patient populations. This meta-analysis integrates data from independent static FDG-PET and fMRI studies across four neurological and psychiatric disorders: Alzheimer’s disease (AD), Parkinson’s disease (PD), major depressive disorder (MDD), and schizophrenia (SCZ), selected based on literature availability. This approach allows us to examine whether regions characterized by pathological metabolic activity also exhibit corresponding disruptions in network topology.

By integrating observations across sensory, cognitive, arousal, and disease states, we reveal a complex landscape of energetic costs underlying fMRI-derived nodal graph-theoretical metrics. Using nodal degree as a primary example, we show that although existing studies have identified a positive association between degree and glucose metabolism at baseline, this relationship is not consistently preserved across brain states. Moreover, its expression is spatially heterogeneous: metabolic and degree changes in the executive control and dorsal and ventral attentional networks show more consistent directional coupling than those observed in other networks. We further demonstrate that this state-dependent relationship with glucose metabolism extends to multiple nodal metrics beyond degree. Finally, we discuss possible mechanisms underlying this complex association, drawing on both theoretical accounts of complex network metrics and empirical evidence for the diverse energy sources supporting brain function. Together, our findings show that nodal graph-theoretical metrics do not carry uniform energetic implications for glucose consumption; rather, their metabolic basis shifts across networks and brain states, necessitating a cautious approach to their interpretation in experimental and clinical settings.

## RESULTS

### Coupling between glucose metabolism and nodal degree varies across brain states: simultaneous fPET-fMRI

We first investigated the association between brain’s regional energy expenditure and nodal degree across 4 brain states: flickering checkerboard-based visual task, finger-tapping motor task, working memory task, and sleep using 4 simultaneous fPET-FDG/fMRI datasets (Fig. 1a). Nodal degree was selected as a primary metric due to its prevalence in graph-theoretical studies and its established positive correlation with glucose metabolism at resting state^13–16^. All analyses were conducted using fPET and fMRI time courses from 300 cortical grey matter ROIs^21^, covering the entire cortical surface. Functional connectivity (FC) matrices were constructed by computing Pearson’s correlation coefficients across all ROI-wise fMRI time courses, resulting in a 300×300 matrix for each brain state. The FC matrices were thresholded by preserving the top 10% strongest positive connections to create binary matrices for calculating the graph theoretical metrics with the Brain Connectivity Toolbox^2^ (see Methods). Task- or sleep-induced changes in nodal degree, relative to rest sessions, were assessed using paired-sample t-tests for each state. Regional task- or sleep-evoked percent signal change (PSC) in CMRglu was calculated from fPET time-activity curves (TACs) using a general linear model (GLM), specifically by comparing the slope of the task portion of the fitted task regressor to the slope of the stable portion of the fitted baseline^22^.

Our glucose metabolism maps for each brain state align with previous literature^17,19,23^: CMRglu increased in primary visual and motor cortices during visual and motor tasks, respectively, increased in frontoparietal and visual regions during working memory task, and decreased globally during non-rapid eye movement (NREM) sleep (Fig. 2, “Glucose metabolism”).

**Figure 2.**
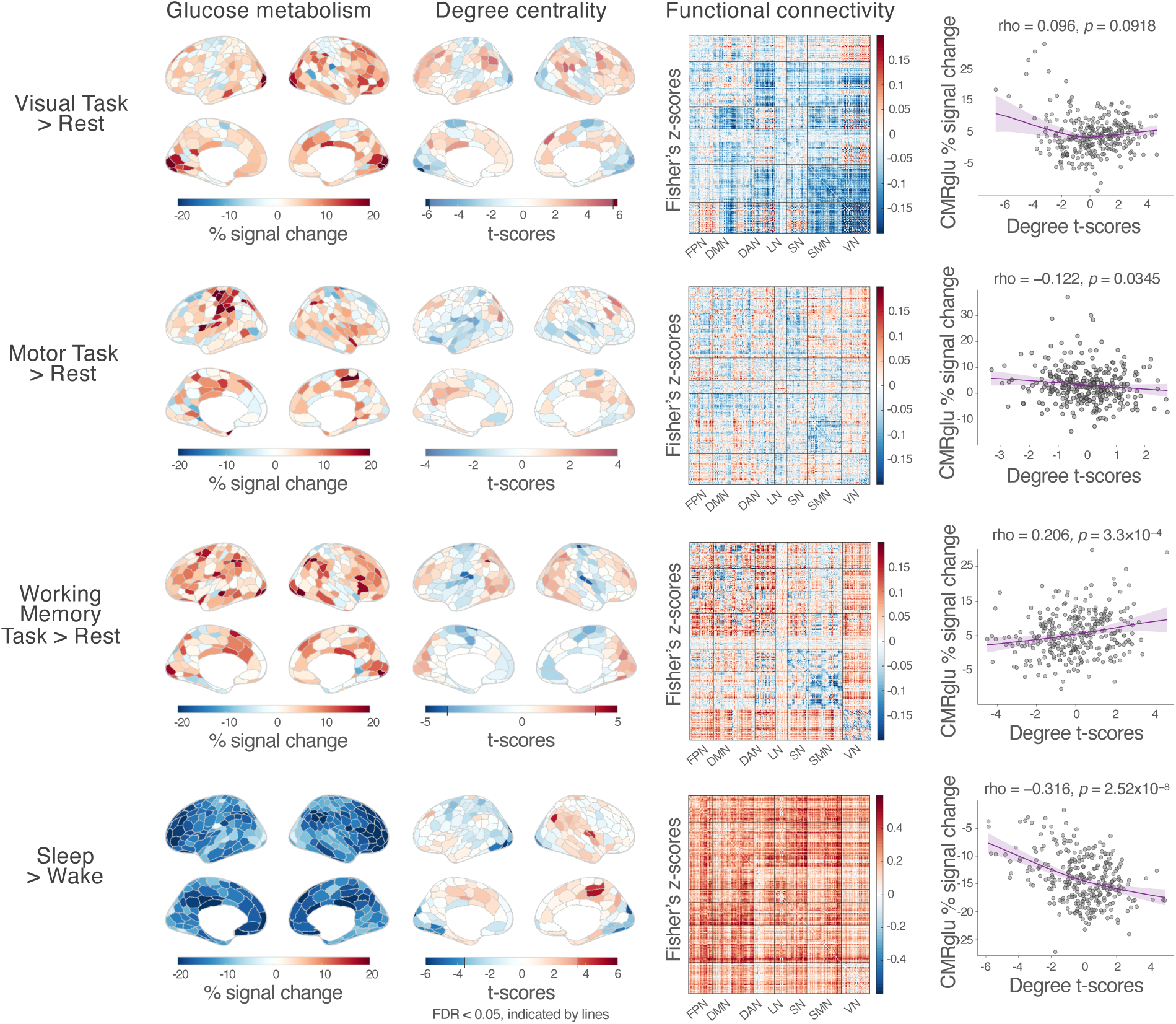
Coupling between regional glucose metabolism and nodal degree changes across sensory, cognitive, and arousal states. Glucose metabolism: state-related changes in percent CMRglu derived from fPET-FDG data. Degree centrality: Paired-sample t tests of fMRI-derived degree differences between tasks or sleep sessions and rest; significant results after FDR (*q* < 0.05) correction are shown in bright color while non-significant patterns are preserved in a less saturated color^72^. State-related changes in functional connectivity matrices are provided as references (“FPN”: frontal parietal network, “DMN”: default mode network, “DAN”: dorsal attentional network, “LN”: limbic network, “SN”: salience network, “SMN”: somatomotor network, “VN”: visual network). Scatterplots depict the relationship between percent CMRglu changes and degree centrality changes for each brain state, with a loess fitted non-linear regression line with 95% confidence interval shaded in purple. Each gray dot represents one cortical ROI. The correlation between t-scores of degree and percent changes in CMRglu was assessed with Spearman’s correlation test.

State-related changes in fMRI-based degree maps present a more complex pattern that diverges from task activation: degree decreased in the visual cortex during visual task, showed no significant changes during motor task, decreased in SMN during working memory task, and decreased in visual cortex but increased in parietal and temporal regions during NREM sleep (Fig. 2, “Degree centrality”). These spatial patterns in degree maps mirrored the complex alterations in FC both between and within networks (Fig. 2, “Functional connectivity”). The FC within the visual network (VN) and between VN and the somatomotor network (SMN) sharply decreased during the visual task compared to rest, driving the decrease of degree in the visual cortex. The FC during motor task had negligible changes compared to rest but revealed a slightly lower FC within SMN. During the working memory task, the FC was characterized by increased FC between VN and all other networks, and between the frontoparietal network (FPN) and dorsal attentional network (DAN), and decreased FC within SMN. Consistent with previous literature^24^, the FC was higher across the whole brain during NREM sleep compared to rest, especially in FPN, DAN and the default mode network (DMN), driven by the more synchronized brain activities during NREM sleep stages^25^.

The relationship between changes in CMRglu and degree during different task/arousal states was assessed through their spatial correlation across the cortex. Percent changes in CMRglu were negatively correlated with t-scores of degree change from rest to motor task (Spearman’s rho = −0.122, *p* = 0.0345) and from wake to NREM sleep (Spearman’s rho = −0.316, *p* = 2.5×10^−8^), while positively correlated with t-scores of degree change from rest to visual (Spearman’s rho = 0.096, *p* = 0.0918) and working memory (Spearman’s rho = 0.206, *p* = 3.3×10^−4^) tasks. These observations revealed variable polarities and strengths of coupling between glucose metabolism and nodal degree across different brain states.

### Coupling between glucose metabolism and nodal degree varies across brain states: meta-analysis of disease

To further investigate the relationship between glucose metabolism and complex network topology across diverse contexts, including clinical settings, we conducted a meta-analysis on existing static FDG-PET and fMRI studies reporting regional changes in glucose uptake or degree in patients with 4 neuropsychiatric diseases (Fig. 1b): AD (n = 18 PET studies, 9 fMRI studies), PD (n = 11 PET studies, 19 fMRI studies), MDD (n = 6 PET studies, 11 fMRI studies), and SCZ (n = 6 PET studies, 10 fMRI studies). Demographic information of included studies is summarized in Table 1.

**Table 1.**
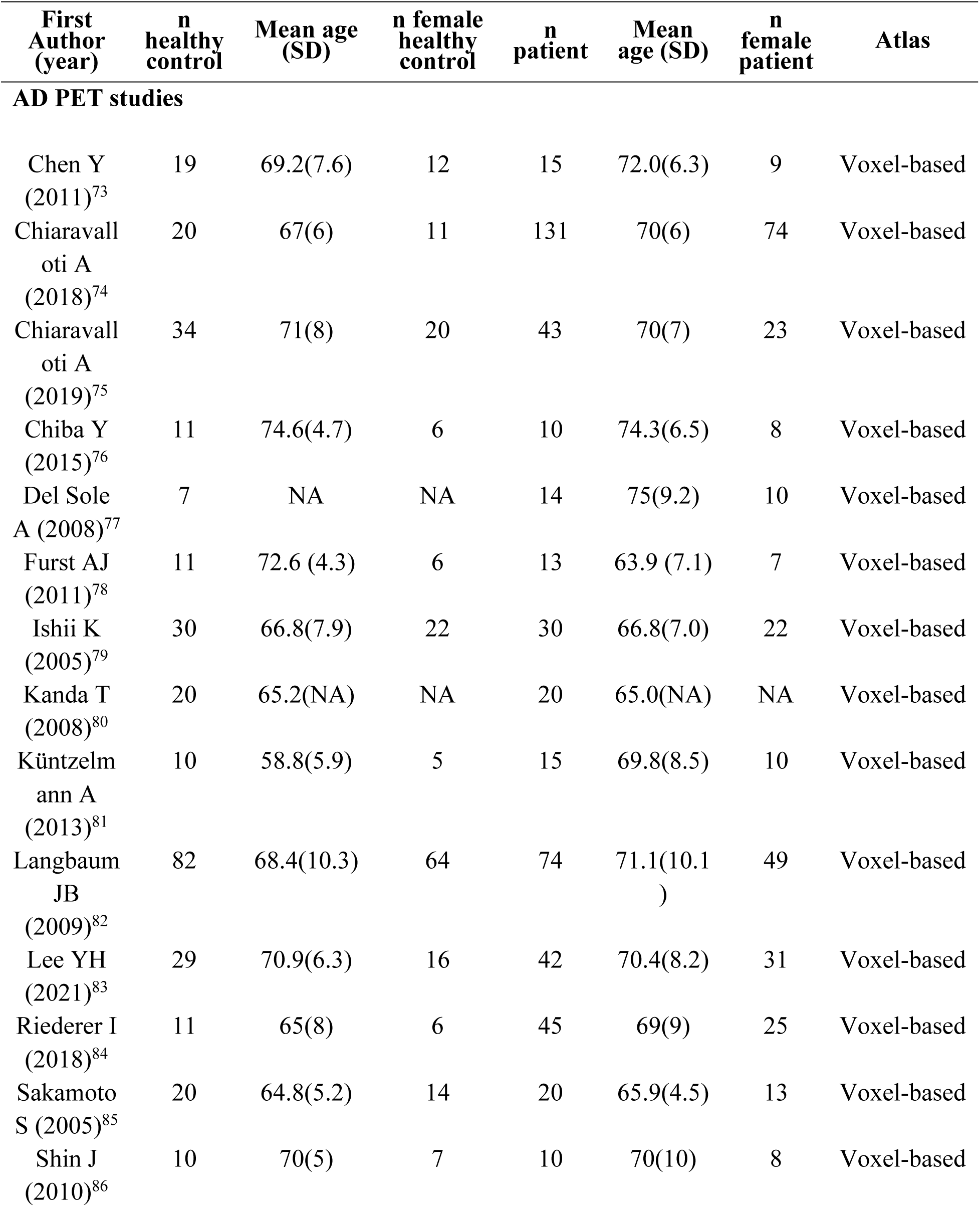
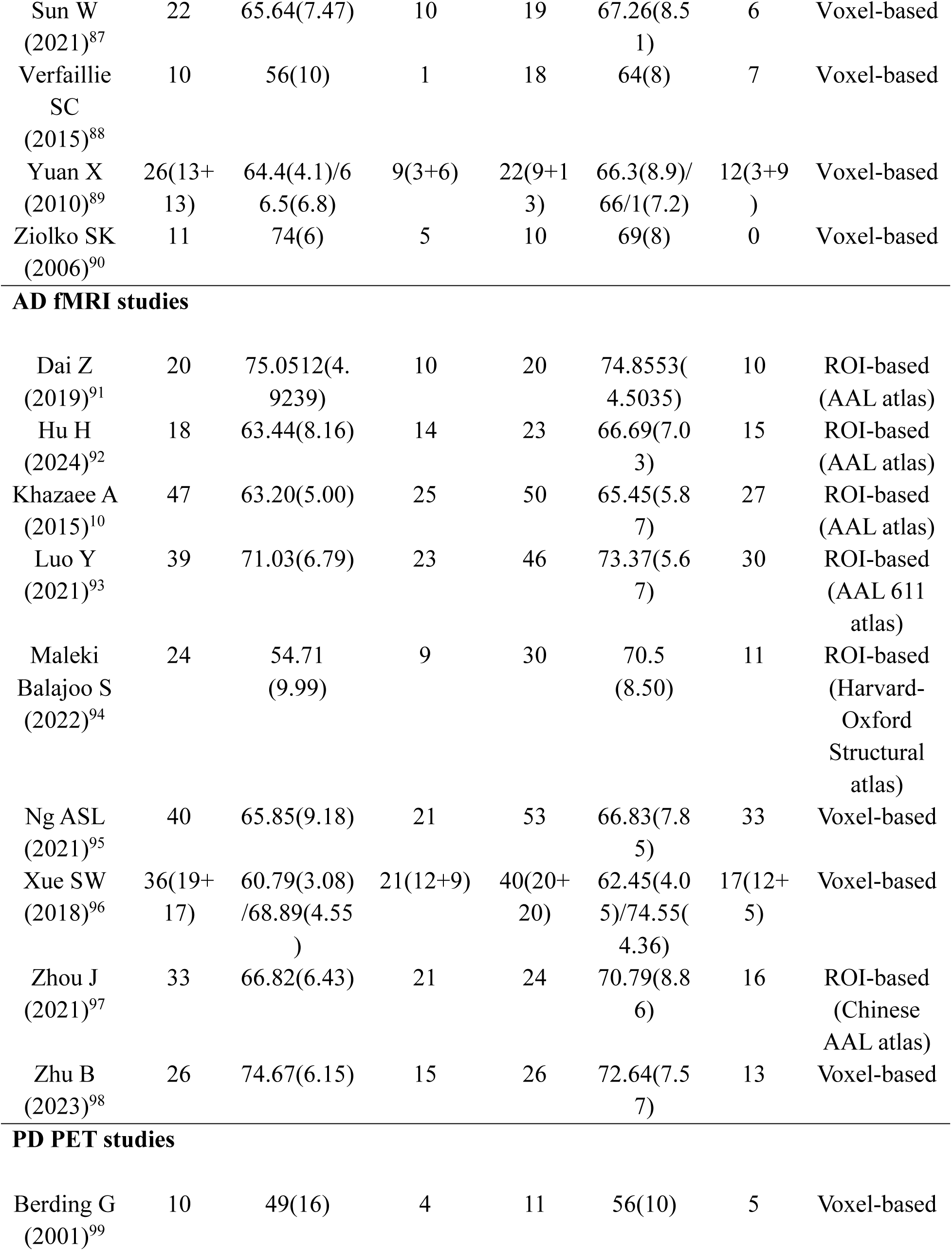
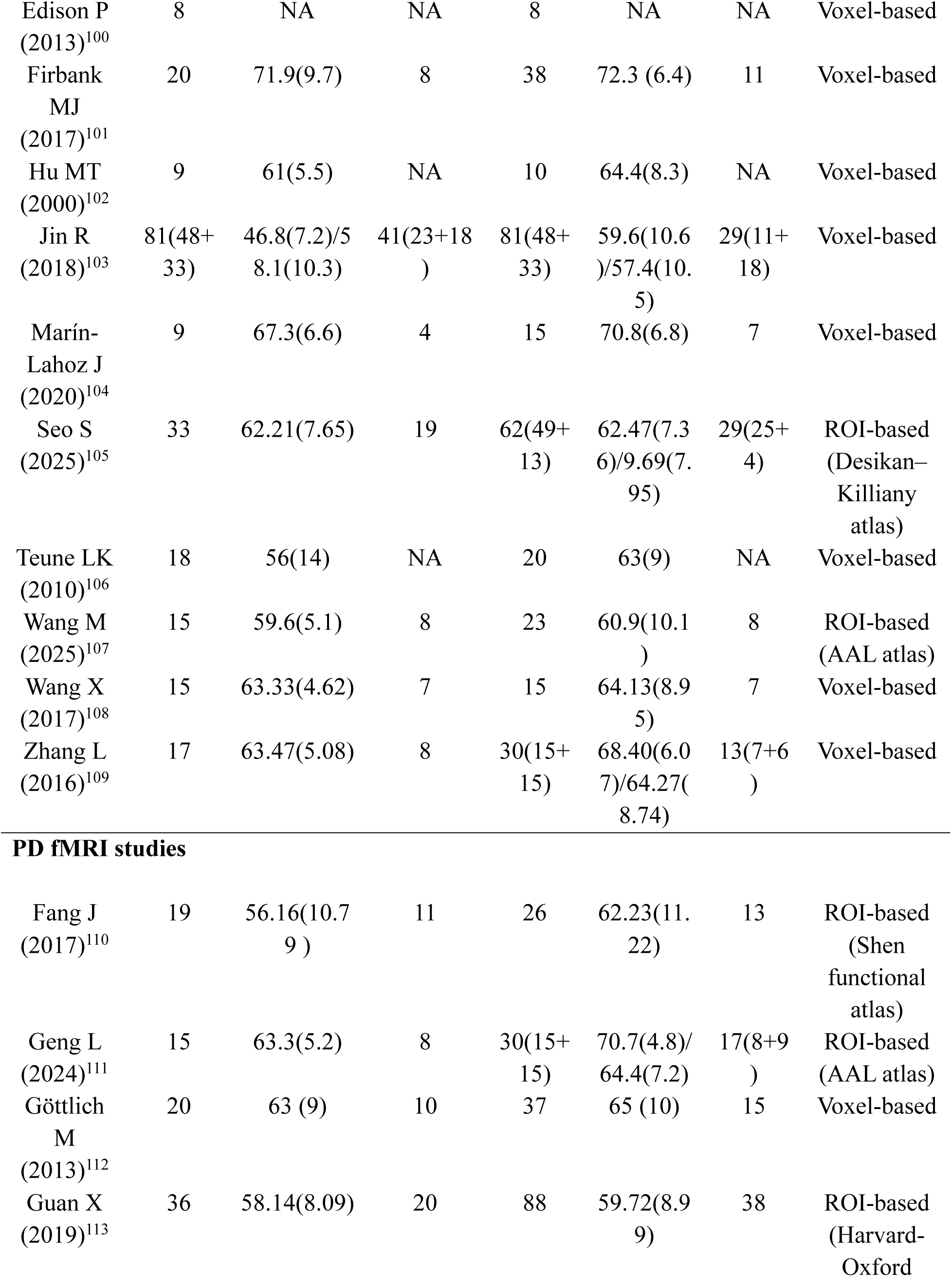
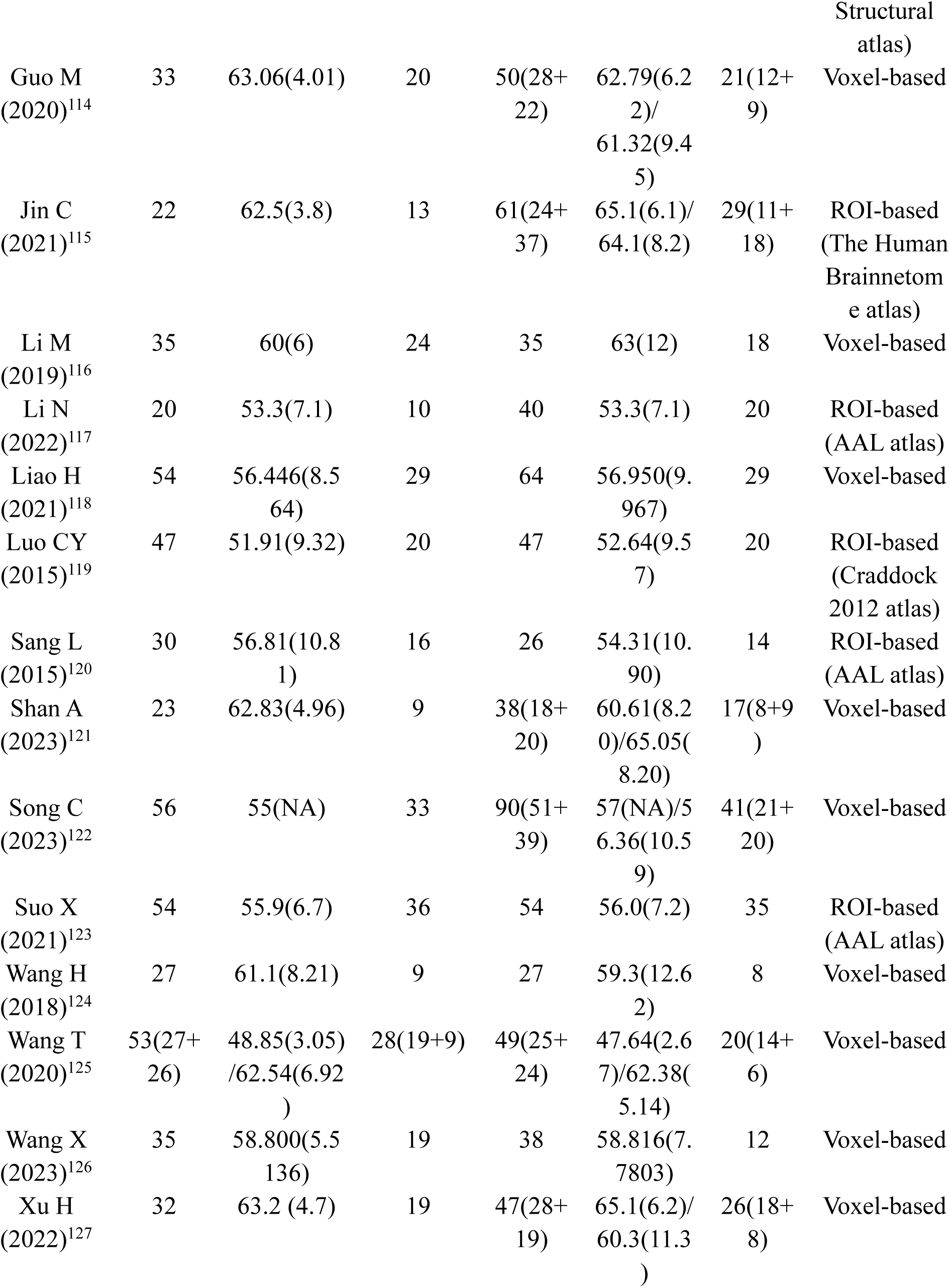
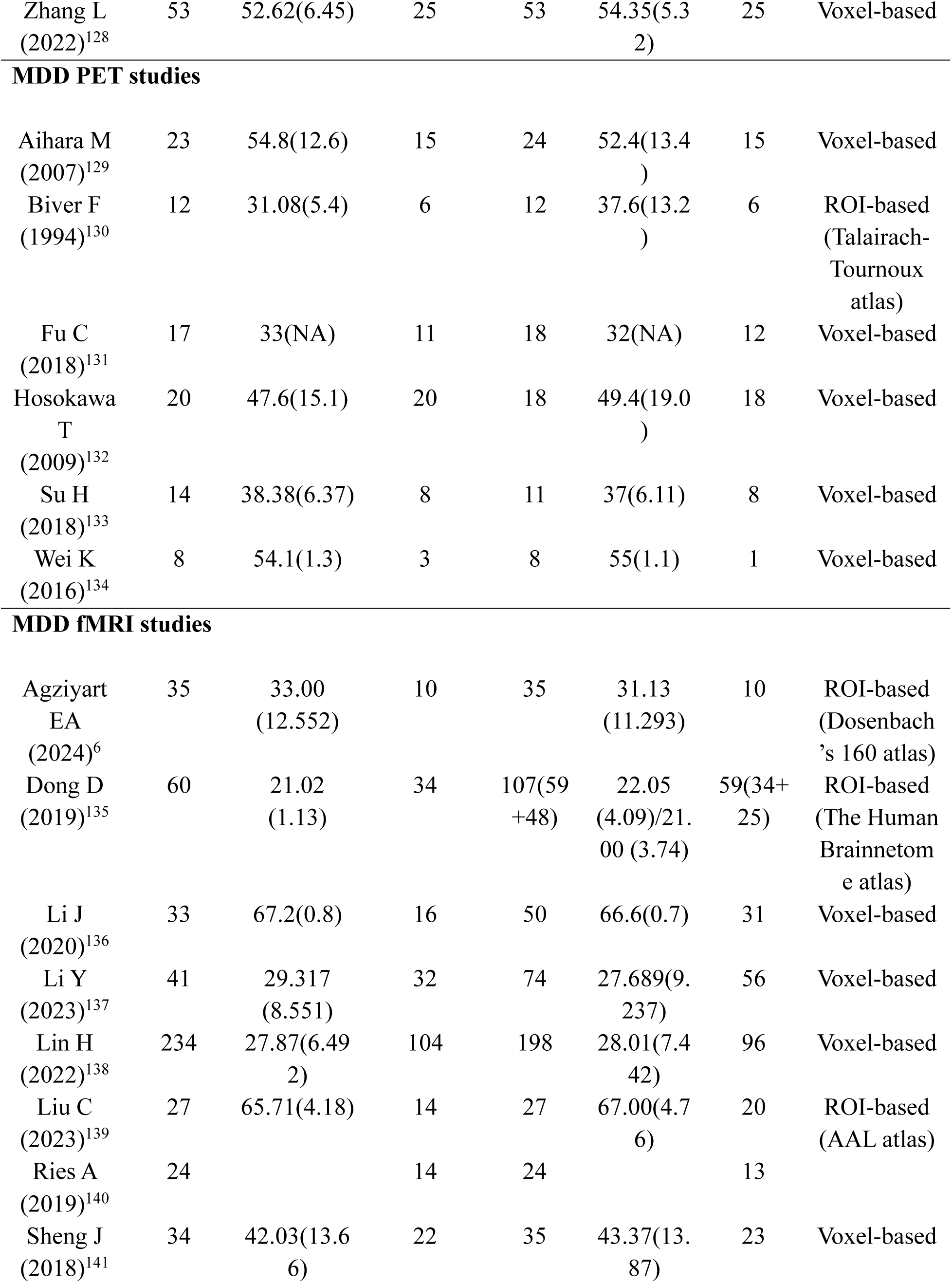
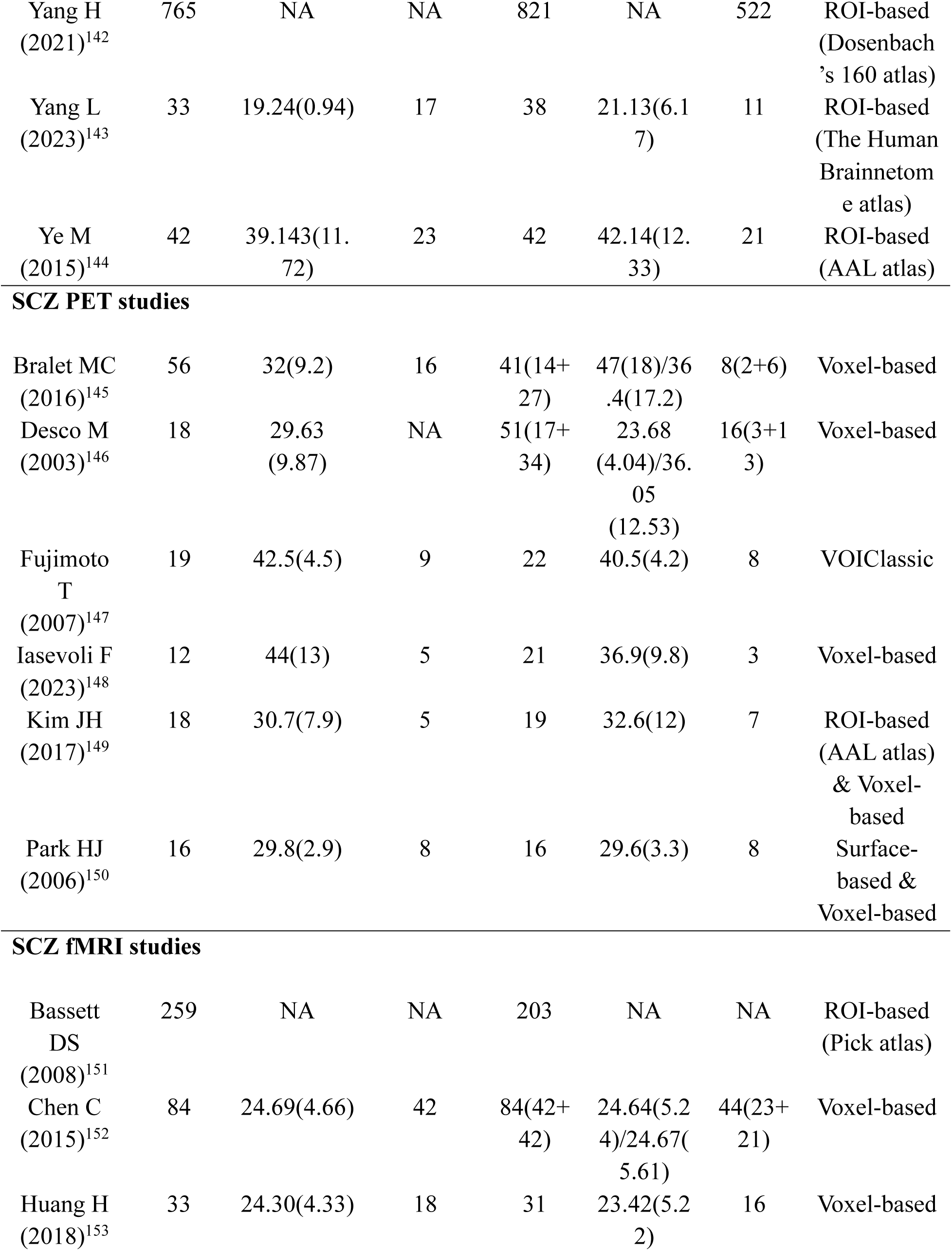
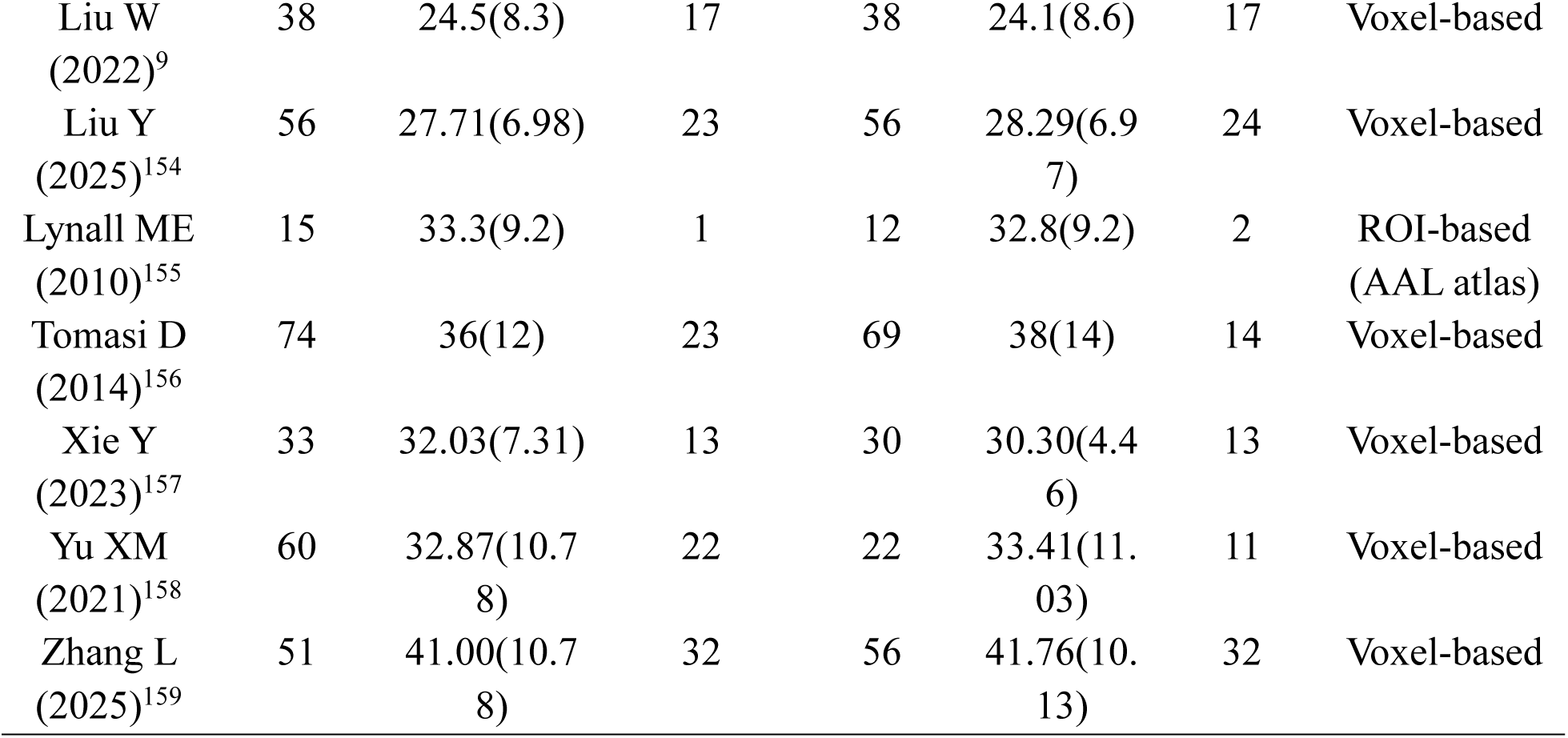
Studies included in the meta-analysis of disease-related changes in regional glucose metabolism and nodal degree.

Owing to the lack of coordinates reported from several earlier studies included in the meta-analysis, we performed the analysis at region of interest (ROI)-level with the most commonly used brain parcellation scheme among all the included studies, the Automated Anatomical Labeling (AAL)^26^ 90-region atlas. We calculated the probability for each region to be reported as having significant changes in glucose uptake or degree in patients compared to healthy controls among identified studies.

The meta-analysis results are summarized in Fig. 3. In AD, we identified robust whole-brain hypometabolism, accompanied by decreased nodal degree in the precuneus, temporal and right medial frontal regions, and increased degree in the left lateral frontal and bilateral parietal regions. In PD, we found hypometabolism in selected occipital, parietal, and temporal regions, lower degree in portions of the occipital and parietal regions, and higher degree in the frontal cortex and precuneus. The results across PD studies were relatively heterogeneous, potentially reflecting the wide range of its clinical symptoms^27^. In MDD, we observed hypometabolism in prefrontal, parietal, and temporal regions, alongside a widespread reduction in nodal degree across the brain with the exception of the right parietal and temporal cortex and limbic regions. In SCZ, we identified consistent hypometabolism in the frontal cortex and hypermetabolism in the temporal cortex. The alignment of degree alteration among SCZ studies is weak and asymmetrical, which may similarly reflect heterogeneity in symptom profiles and clinical subtypes within SCZ populations.

**Figure 3.**
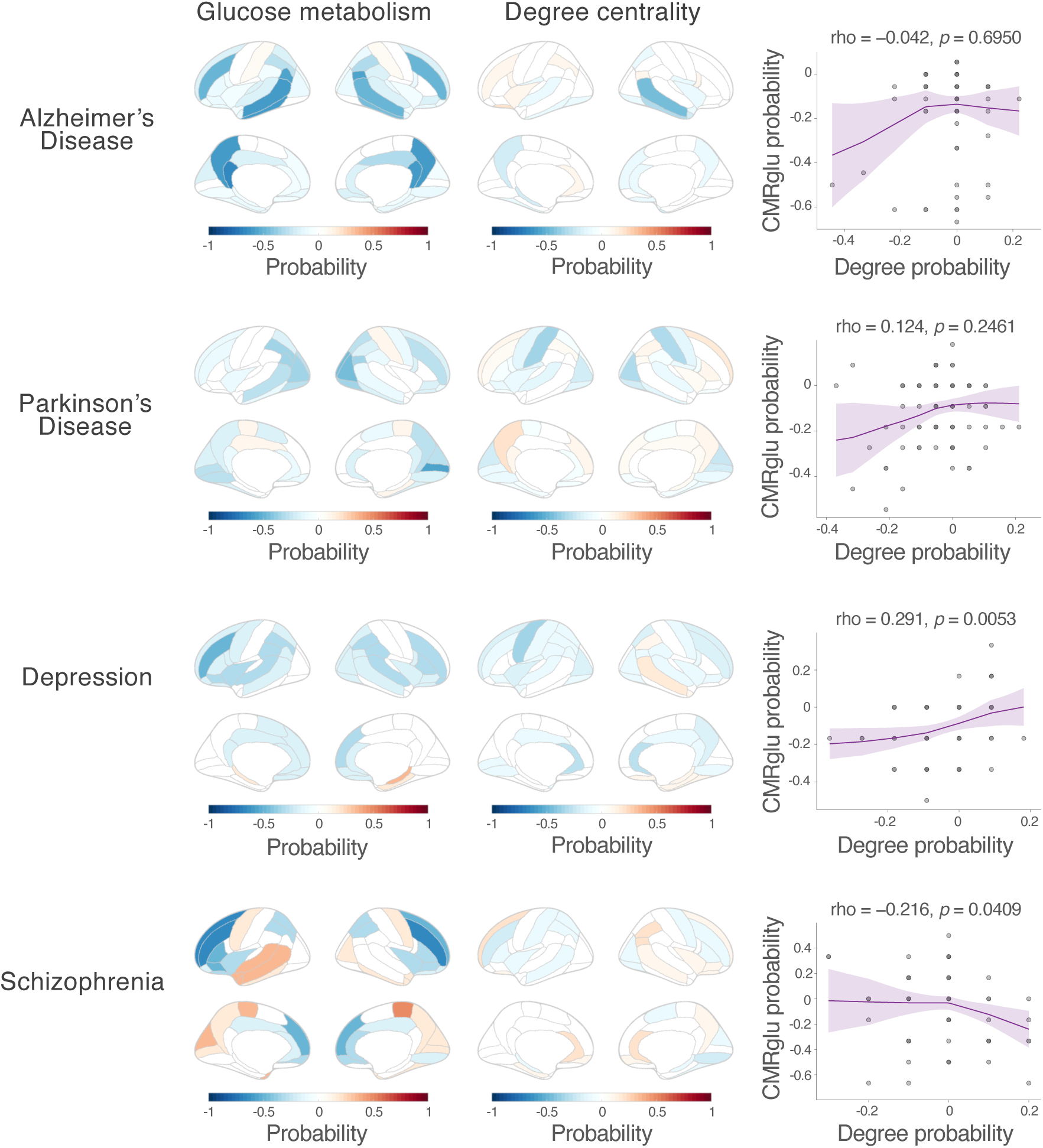
Coupling between regional glucose metabolism and nodal degree changes across diseases derived from meta-analysis. The “likelihood of activation” for each brain region is estimated as the probability that the region is reported to exhibit significant alterations across all included studies for each ROI in the AAL 90-ROI atlas. Red (positive) indicates increased glucose metabolism or degree centrality in patients compared to healthy subjects, and blue (negative) indicates decreases. The scatterplot depicts the relationship between the likelihood of activation for glucose metabolism and nodal degree changes for each disease, with a loess fitted regression line with 95% confidence interval shaded in purple. Each gray dot represents one brain ROI. The correlation between activation probability of degree and glucose metabolism was assessed with Spearman’s correlation test.

The alignment between disease-related alterations in glucose uptake and nodal degree was quantified by computing the spatial correlation of their ROI-wise activation likelihood maps for each disease. We found a positive trend in PD patients (Spearman’s rho = 0.124, *p* = 0.2461), a significantly positive correlation in MDD patients (Spearman’s rho = 0.291, *p* = 0.0053), and a negative trend in AD patients (Spearman’s rho = −0.042, *p* = 0.6950) and SCZ patients (Spearman’s rho = −0.216, *p* = 0.0409). Together, these observations indicate the absence of a consistent global relationship between glucose metabolism and nodal degree across disease states, in line with patterns observed in functional PET-MRI data across task and arousal states.

### Network-specific coupling between glucose metabolism and degree

Having identified a state-dependent, incongruent association between glucose metabolism and nodal degree at the global level, we further evaluated whether this complex relationship manifests consistently across different functional brain networks. Using the Yeo 7-Network parcellation^28^, we characterized the network-specific consistency in the polarity of metabolic and nodal degree changes across different brain states. To quantify this, we generated an index map for each state, assigning a value of 1 to regions where glucose metabolism and degree changed in the same direction, and –1 where they diverged. These maps were then averaged across states to produce consistency maps.

We found that across task and arousal states, regions in FPN and DAN showed more consistent directional coupling between glucose metabolism and degree; across diseases, regions within the FPN, DAN and the salience network (SN) exhibited more consistent directional coupling between glucose metabolism and degree (Fig. 4). These observations suggest that the coupling between nodal degree and glucose metabolism varies spatially, with altered nodal degrees in transmodal, higher-order control networks more likely reflecting changes in underlying metabolic activity.

**Figure 4.**
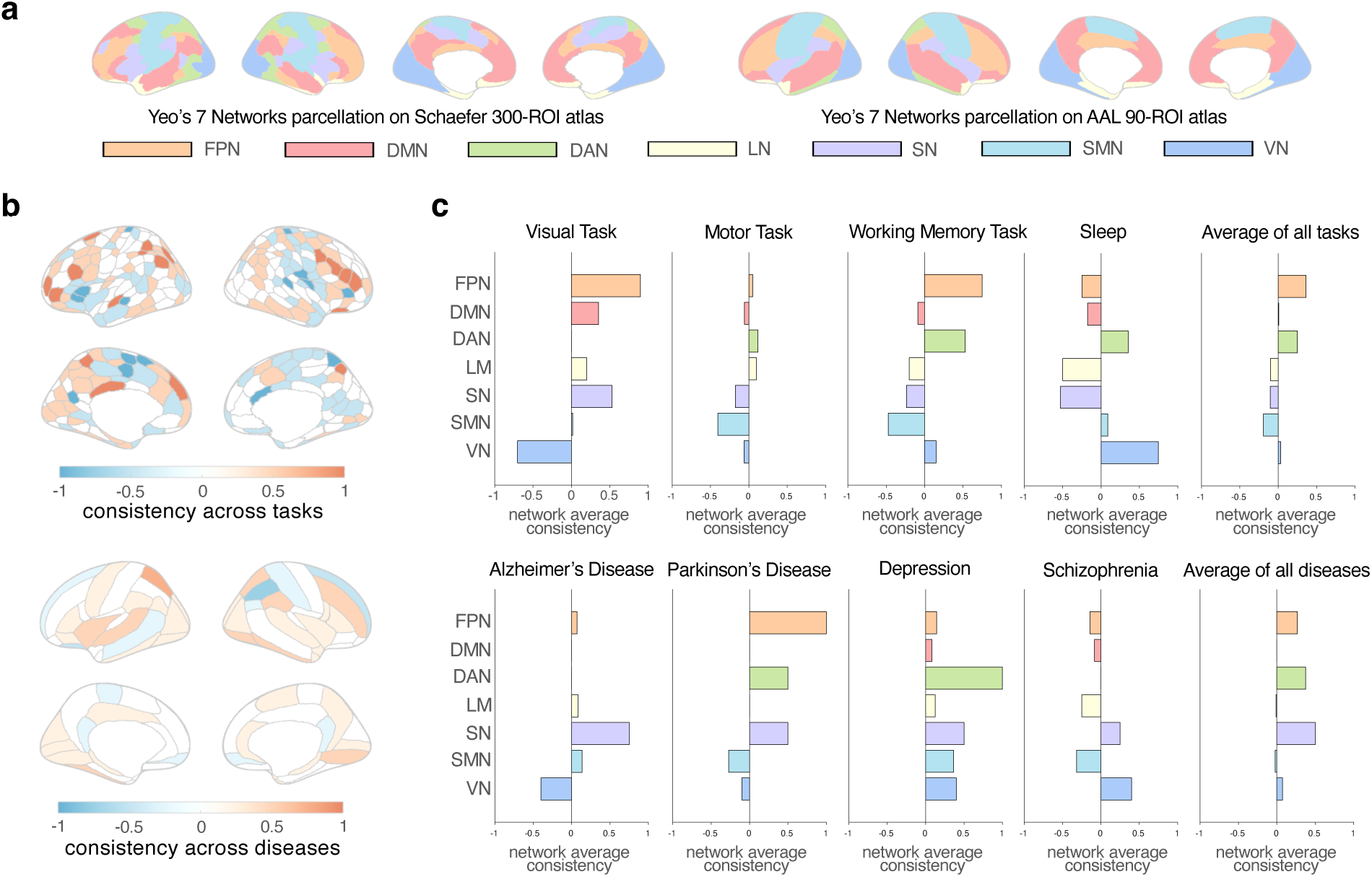
Network-specific coupling between regional glucose metabolism and nodal degree across brain states. (a) Network parcellation scheme used to derive network-specific coupling patterns based on Yeo’s 7 Networks parcellation projected onto the Schaefer 300-ROI and AAL 90-ROI atlases. (b) Regional consistency in the polarity of changes in glucose metabolism and nodal degree, averaged across four task- or sleep-evoked brain states and four disease-related brain states, to assess the stability of (de)coupling across states. Red (positive) indicates a consistent direction of change in glucose metabolism and nodal degree, whereas blue (negative) indicates inconsistency. (c) Network-level consistency in the polarity of changes in glucose metabolism and nodal degree, calculated by averaging the consistency value across all regions within each brain network.

### Metabolic correlates of nodal graph-theoretical metrics beyond degree

Graph theory offers a wide range of metrics that characterize different properties of networks, and nodal centrality assessed by degree is only one facet. We further assessed whether other graph-theoretical metrics, beyond degree, demonstrate consistent relevance to brain energy expenditure across different brain states. Specifically, we linked glucose metabolism to another five common FC-based nodal graph-theoretical metrics for the task and sleep datasets: clustering coefficient, local efficiency, nodal efficiency, within-module degree z-score, and participation coefficient (Fig. 5a; see Methods for mathematical definitions). This analysis was restricted to the empirical fPET-fMRI data due to fewer publications on graph-theoretical metrics beyond nodal degree suitable for meta-analysis. Task-related changes in these metrics are shown in Supplementary Fig. 1.

**Figure 5.**
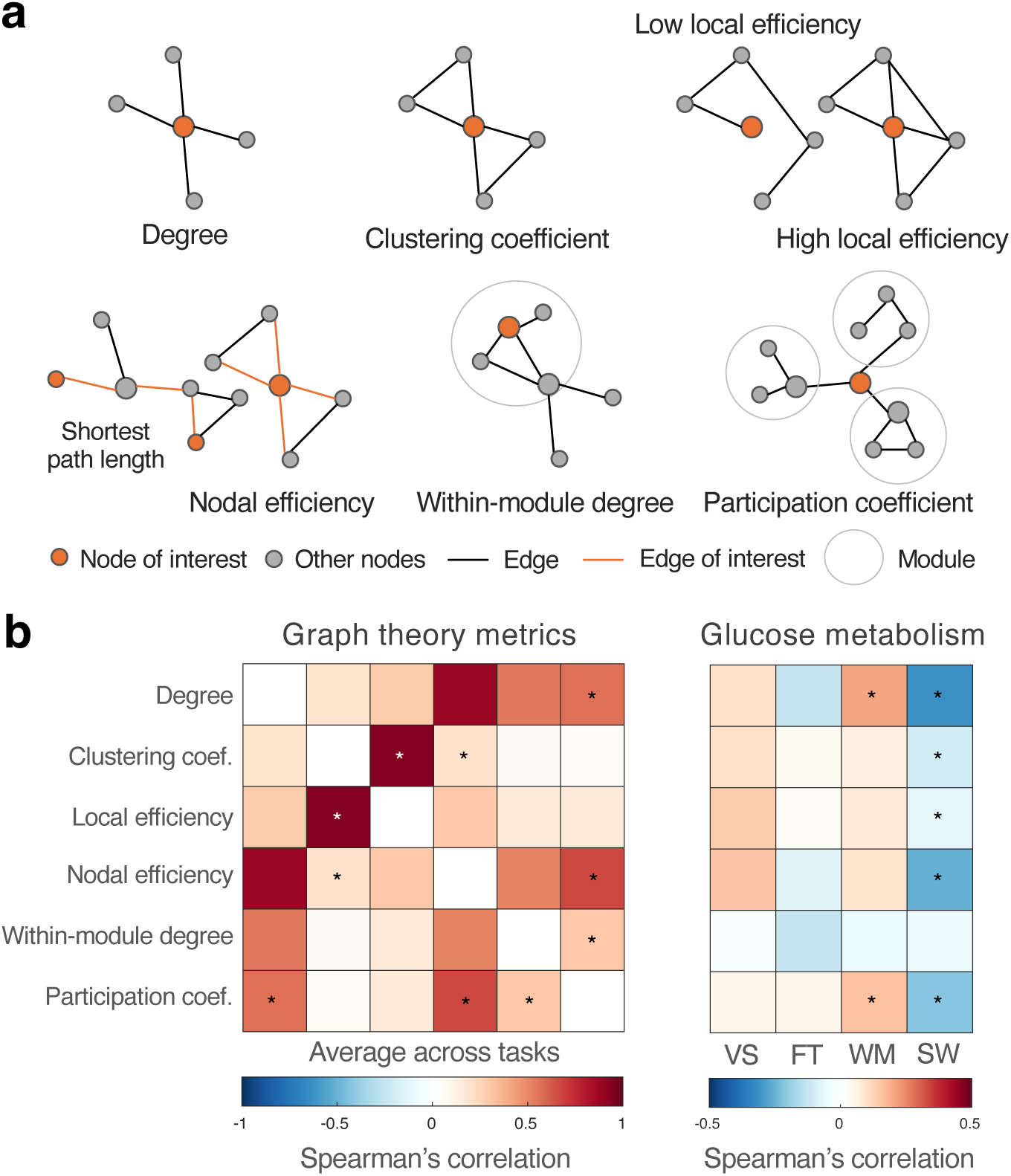
Coupling of regional glucose metabolism and nodal graph-theoretical metrics beyond degree. (a) Cartoon illustration of different graph-theory metrics examined in this study. Each dot represents a node/brain region, and each line represents an edge/functional connection between brain regions. Orange dot/line represents node/edge of interest for each metric, e.g., the orange dot in the first graph has high degree. The grey circles represent network modules. Degree quantifies the number of connections a node has; clustering coefficient measures how tightly interconnected a node’s neighbors are; local efficiency reflects how efficiently its neighbors can communicate when a node is removed; nodal efficiency measures how efficiently a node integrates information across the whole graph by having short path length to other nodes; within-module degree quantifies intra-modular connectivity; participation coefficient reflects inter-modular connectivity. (b) Correlations amongst graph-theoretical metrics (left, averaged across all brain states) and between glucose metabolism and graph-theoretical metrics (right, for each brain state), estimated using Spearman’s spatial correlation test and simultaneous fPET-fMRI datasets across task and sleep states. Red indicates a positive correlation, and blue indicates a negative correlation. “VS” = visual task, “FT” = motor (finger-tapping) task, “WM” = working memory task, “SW” = sleep (sleep-wake transitions). Asterisks indicate correlations exceeding the 97.5th or below the 2.5th percentile of a null distribution (5000 permutations).

Correlation between all six complex network metrics (including degree) was assessed with Spearman’s correlation test and averaged across tasks, and their correlation with glucose uptake was assessed for each task (Fig. 5b). The correlation structure among different network metrics was highly consistent across tasks, likely reflecting both their interrelated definitions and their shared dependence on the underlying spatiotemporal structure of the imaging data^29^: degree and nodal efficiency were strongly coupled, as the latter is derived from degree; within-module degree *z*-score and participation coefficient were also correlated with degree, as they reflect within- and across-module centrality; and clustering coefficient and local efficiency were tightly coupled together, as they both index local integration. However, like nodal degree, the metabolic correlates of other nodal metrics varied across tasks. All nodal metrics, except for the within-module degree z-score, had both positive and negative correlations with glucose metabolism across tasks. These observations suggest that the dynamic, state-dependent relationship with cerebral energetic costs is not unique to degree metrics, but rather extends to a broad range of nodal graph-theoretical measures commonly used to characterize large-scale brain network organization.

## DISCUSSION

By integrating empirical functional PET-MRI with meta-analytical approaches, we show that commonly used nodal graph-theoretical metrics do not carry consistent implications for altered cerebral glucose metabolism, revealing a dissociation between complex network topology and brain metabolic activity across diverse brain states. In particular, the presumed monotonic coupling between energy expenditure and nodal degree, often interpreted as higher metabolic demand associated with denser functional connections, does not generalize beyond resting-state conditions, but is instead confined to specific networks supporting integrative functions. The observed complexity of energy-topology coupling likely reflects multiple contributing factors, including state-dependent functional connectivity profiles, the definitions and computational properties of graph-theoretical metrics, and the specific contrast mechanism of fPET-FDG, as discussed below.

Identifying the mechanistic basis for these diverse metabolism-topology coupling patterns is key to developing accurate neuroenergetic interpretations of graph-theoretical metrics. Figure 6 offers a conceptual framework for understanding why functional network topology can deviate from underlying metabolic costs, using a few illustrative scenarios centered on nodal degree and assuming a close coupling between neural activity and metabolism. This figure, specifically, outlines a few conditions under which increased neural activity and metabolism may be associated with opposite (a, b), unchanged (c), or concordant (d, e) changes in nodal degree. These theoretical scenarios recapitulate the metabolism-degree coupling patterns observed in our data.

**Figure 6.**
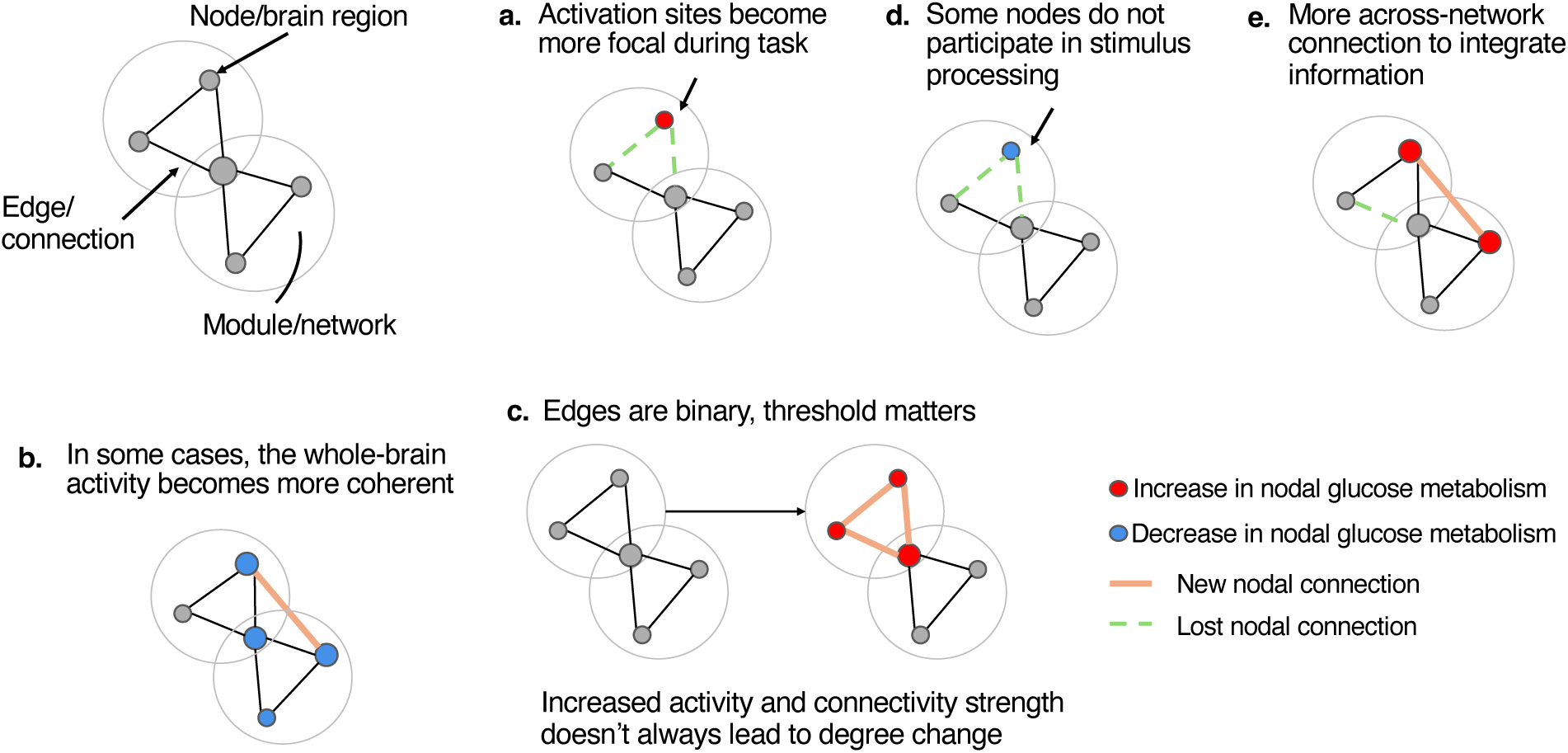
Cartoon illustration of the complex, state-dependent coupling between glucose metabolism and nodal degree. (a) Theoretical account for complex couplings between glucose metabolism and nodal degree. Red dot indicates increases in glucose metabolism, and blue dot indicates decreases; thickened orange line indicates new connection between brain regions, and dashed green line indicates lost connection. (b) Spatial correlation between degree and different metabolic pathways. Degree spatial maps (magenta) and CMRglu (orange) were quantified from a publicly available resting-state fPET-fMRI dataset^69^. The spatial correlation between resting-state degree and different metabolic pathways (including CMRglu) was assessed with Spearman’s correlation test. The spatial maps of different metabolic pathways were extracted from a public dataset^55^. Degree is strongly correlated with energy metabolism pathways of PPP (Spearman’s rho = .4370, *p* = 4.36×10^−20^), TCA (Spearman’s rho = 0.3009, *p* = 8.12×10^−10^), and lactate (Spearman’s rho = 0.2576, *p* = 5.33×10^−7^), and not significantly correlated with glycolysis (Spearman’s rho = 0.0231, *p* = 0.6444), oxidative phosphorylation (Spearman’s rho = −0.0102, *p* = 0.8395), nor CMRglu (Spearman’s rho = 0.0306, *p* = 0.5424). Significant correlations after FDR correction (*q* < 0.05) are marked with an asterisk.

**Figure 7.**
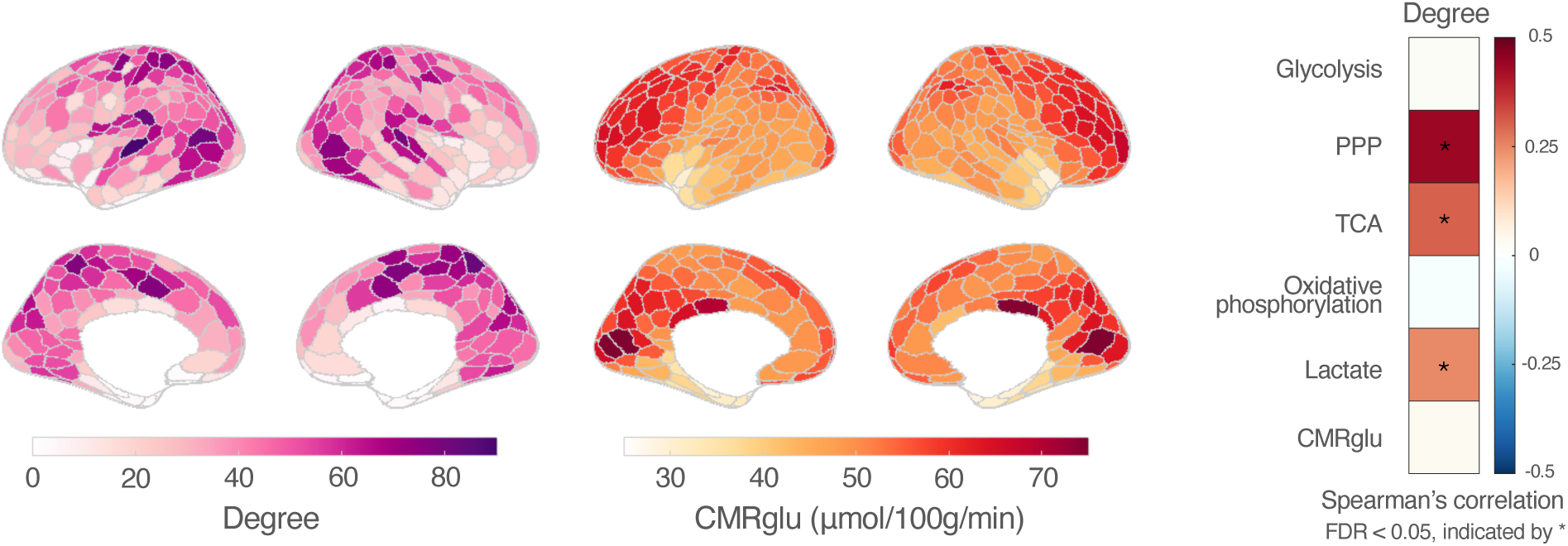
Spatial correlation between resting-state degree and different metabolic pathways. Degree spatial maps (magenta) and CMRglu (orange) were quantified from a publicly available resting-state fPET-fMRI dataset^69^. The spatial correlation between resting-state degree and different metabolic pathways (including CMRglu) was assessed with Spearman’s correlation test. The spatial maps of different metabolic pathways were extracted from a public dataset^55^. Degree is strongly correlated with energy metabolism pathways of PPP (Spearman’s rho = .4370, *p* = 4.36×10^−20^), TCA (Spearman’s rho = 0.3009, *p* = 8.12×10^−10^), and lactate (Spearman’s rho = 0.2576, *p* = 5.33×10^−7^), and not significantly correlated with glycolysis (Spearman’s rho = 0.0231, *p* = 0.6444), oxidative phosphorylation (Spearman’s rho = −0.0102, *p* = 0.8395), nor CMRglu (Spearman’s rho = 0.0306, *p* = 0.5424). Significant correlations after FDR correction (*q* < 0.05) are marked with an asterisk.

For instance, in the sensory task example (Fig. 2, “Visual Task > Rest”), while increased CMRglu in visual cortex reflects task-evoked neuronal activation, more focal activations induced by unchanging sensory stimulation/task (e.g. flickering checkerboard)^30^ can result in decreased functional connections and therefore degree (Fig. 6a). Prior studies have reported decreased connectivity within VN^31,32^ and between visual cortex and higher-order visual regions^33^ during visual tasks compared to rest. This observation is consistent with models of the visual system as a hierarchical prediction engine^34^, where predictable, unchanging stimuli minimize prediction error and subsequently lower FC within VN^32^. A similar pattern is observed in the motor task condition (Fig. 2, “Motor Task > Rest”) where we find increased CMRglu alongside decreased FC within SMN. This result is consistent with previous reports of unilateral finger tapping results in anti-correlation of bilateral motor cortices^32,35,36^, suggesting lower coherence driven by focal activations.

The decoupling of CMRglu and degree during sleep is better captured by Fig. 6b. During early NREM sleep, neuronal activity becomes more coherent and synchronized, leading to increased FC strength across cortical regions (Fig. 2, “Sleep > Wake”). Yet this heightened connectivity coincides with reduced glucose consumption, most prominently in the DMN, a network that is metabolically and functionally active during wakeful rest, supporting self-referential thought and conscious awareness^37^. This dissociation illustrates how increased network connectivity does not necessarily reflect increased metabolic demand.

Beyond neuronal mechanisms, the apparent dissociation of metabolism-degree coupling may also arise from the methodological constraints of graph construction. Specifically, during the thresholding and binarization of FC matrices, information regarding connection strength is discarded, rendering nodal degree estimates highly sensitive to the chosen threshold (Fig. 6c). This, for example, explains our observation that while sleep significantly increased global FC strength, the resulting spatial map of nodal degree changes had a mixture of positive and negative t-scores (Fig. 2, “Sleep > Wake”). In our primary analysis, which preserved the top 10% of connections of FC matrices, the degree changes primarily reflected changes in nodal hubness driven by network reorganization rather than absolute changes in FC strength.

To evaluate the sensitivity of results on binarization thresholds, we further assessed the consistency of network metric changes across tasks and arousal states using different thresholds. Degree, nodal efficiency, within-module degree, and participation coefficient were relatively stable and highly positively correlated across thresholds, whereas clustering coefficient and local efficiency showed variability and moderate positive correlation across thresholds (Supplemental Figs. 2). Although nodal degrees calculated at different thresholds are highly correlated, these thresholds modulate the coupling between nodal degree and glucose metabolism, and their modulatory effect also varies across states (Supplementary Fig. 3). Despite having similar spatial patterns across thresholds, nodal metrics calculated from different thresholds favor connections of different strengths: higher thresholds (e.g., top 5%) preserve only the strongest connections, while lower thresholds (e.g., top 40% or a weighted network) include progressively weaker connections. If the (de)coupling between glucose metabolism and nodal degree is driven by specific regions exhibiting strong alignment or misalignment, including weaker connections may dilute their impact on the global spatial correlation. This methodological sensitivity cautions against assuming a fixed relationship between network topology and metabolic activity, as both brain state and analytical choices affect the strength and direction of their coupling.

Despite the opposing metabolism-degree examples discussed above, concordant changes in glucose metabolism and nodal degree can occur when task-irrelevant regions reduce both connectivity and metabolic demand (Figure 6d). For example, regions in SMN have been reported to have lower connectivity during a visual semantic decision-making task^38^, and we observed parallel decreases in both glucose metabolism and degree in SMN during the visual task, reflecting reallocation of metabolic and network resources to support task-relevant activity.

Metabolic activity and nodal degree can also show synchronized activity when energy demands are primarily driven by the need for information integration. In this case, the metabolic cost of a node is intrinsically tied to its connectivity with other brain regions, a mechanism similar to the monotonic relationship observed in the resting state^13–16^. Specifically, when a node maintains an integrative hub role across various brain states (Fig. 6e), its metabolic activity more tightly tracks its topological profiles.

This relationship may explain why specific networks, such as the FPN and DAN, are more likely to have aligned changes in glucose metabolism and nodal degree across task and disease states (Fig. 3c). A large body of literature characterizes the FPN as a collection of flexible hubs that integrate and mediate across-network activity to accommodate task demands and cognitive control^12,39–42^, with a prominent role in regulating the switch between DMN and DAN to suppress self-referential thought during goal-directed tasks^39,43^. Supporting this, we observed increased glucose metabolism in FPN across sensory and cognitive tasks, accompanied by a higher degree and participation coefficient, an index of participation in network integration, among the same or adjacent regions (Supplementary Fig. 1). Similarly, DAN collaborates with sensory networks, especially VN due to the visual nature of most fMRI tasks^41,44^, to facilitate top-down or stimulus-driven attentional control^45^. Thus, the stronger coupling between glucose metabolism and degree in the FPN and DAN likely reflects the elevated energetic demands of task-evoked cross-network regulation and integrative processing.

This framework also provides a plausible explanation for the coupled metabolism-degree relationship observed in FPN and DAN in clinical populations. Because these hub regions support multiple core cognitive functions and require more energy to maintain their complex interactive roles^46^, they are more susceptible to a wide range of disease-related stressors^8,47^. For example, cerebral hypometabolism in diseases like AD, PD, MDD and SCZ may arise from gray-matter atrophy^48^, oxidative stress^49,50^, insulin resistance^51–53^, and dopamine deficits^53,54^. When energetic supply is insufficient to sustain extensive connectivity, or when neuronal loss directly disrupts network architecture, these hub regions are unable to maintain a high number of functional connections, leading to concomitant reductions in nodal degree.

Finally, while we focused on glucose metabolism captured by FDG-PET, brain energy metabolism encompasses diverse biochemical pathways that collectively support neural function. More definitive associations between the energetic costs of complex network topology and underlying physiology may require consideration of this broader metabolic landscape. In particular, cerebral glucose metabolism alone comprises several parallel pathways, including glycolysis followed by oxidative phosphorylation, driven by pyruvate entering the tricarboxylic acid (TCA) cycle and fueled by lactate, pentose phosphate pathway (PPP), and glycogen synthase^55^. FDG-based measurements primarily reflect the initial glucose uptake, as FDG cannot be further metabolized once phosphorylated by hexokinase to FDG-6 phosphate^56^. While FDG-based estimates of CMRglu have therefore served as the principal metabolic measure in most prior studies^57^, largely owing to their accessibility, our preliminary analysis revealed stronger resting-state correlations between nodal degree and metabolic pathways of PPP, TCA, and lactate than with FDG-based CMRglu. This implies a possibility that, at rest, regions with high nodal hubness may rely on auxiliary metabolic pathways beyond initial glycolysis to sustain ongoing brain activity. Consistent with this interpretation, previous studies have shown that neurons draw substantially on the PPP to generate NADPH for oxidative stress defense, while supplying pyruvate and lactate to the TCA cycle for ATP production, showing a partial uncoupling from glycolysis^58^. These observations support the notion that resting-state FC is maintained by a metabolically diverse ensemble of biochemical pathways that go beyond simple glucose utilization. How these relationships manifest across different brain states remains an open question, but they underscore the intricate metabolic landscape that supports large-scale brain circuitry and the challenges of deriving coherent interpretations about brain energetics from complex network metrics.

In conclusion, our findings show that nodal graph-theoretical metrics exhibit variable metabolic associations across networks and brain states. These findings emphasize the dynamic, state-dependent nature of metabolism-topology interactions, underscoring the need for careful interpretation of neuroenergetic underpinnings of nodal graph-theoretical metrics in experimental and clinical research.

## METHODS

### Functional PET-MRI acquisition and task paradigm

To assess how brain state states mediate the relationship between nodal graph-theoretical metrics and glucose metabolism, we employed four simultaneous fPET-FDG/fMRI datasets acquired under four different states: visual, motor, working memory, and sleep (Fig. 1a).

#### Visual Task Dataset

A publicly available fPET-fMRI dataset of 10 healthy participants (aged 18–48 years, mean = 29.2 years, 9 females) by research groups at Monash University^59^ was analyzed here. The experiments were conducted on a Siemens (Erlangen, Germany) Biograph 3T mMR scanner. Participants underwent a 90-minute PET-MRI scan with flickering checkerboard visual stimulus blocks and eyes-closed resting state blocks in between. T_2_*-weighted echo planar images (EPIs) were acquired during stimulus and resting-state blocks (repetition time (TR) = 2450 ms, echo time (TE) = 30 ms, voxel size = 3 mm isotropic, 44 slices). FDG tracer was administered through constant infusion at a rate of 36 mL/hr. PET data was acquired in list-mode with the ultrashort TE (UTE) MRI for attenuation correction and have a voxel size of 1.39×1.39×2 mm^3^ and a nominal temporal resolution of 1 minute. T_1_-weighted magnetization-prepared rapid gradient echo (MPRAGE) data was collected for anatomical reference and across-modal registration (TR = 1640 ms, TE = 2.34 ms, inversion time (TI) = 900 ms, voxel size = 1 mm isotropic, 176 slices, flip angle = 8°).

The remaining three datasets were collected locally at the Athinoula A. Martinos Center for Biomedical Imaging at Massachusetts General Hospital (MGH). All experiments were conducted on a Siemens 3T Tim MAGNETOM Trio MR scanner with a BrainPET insert. The Institutional Review Board of MGH approved the studies, and written informed consent was provided by all participants. For the motor task dataset, FDG tracer was administered using a constant-infusion protocol^18^. For the working memory task and sleep dataset, FDG tracer was administered using a bolus + constant infusion protocol^18^, and the total dose for each participant was ~10 mCi (bolus 1.7 mCi, constant infusion 8.3 mCi). PET data for all 3 local datasets were sorted into line-of-response space and compressed into signogram space for reconstruction with a standard 3D ordinary Poisson ordered-subset expectation maximization (OP-OSEM) algorithm, during which attenuation correction was performed using a pseudo-CT map constructed with high-resolution structural MRI data^60^. The reconstructed fPET data have a voxel size of 2.5×2.5×2.5 mm^3^ and a nominal temporal resolution of 30 seconds. The MRI acquisition protocols for the three datasets are described below.

#### Motor Task Dataset

Seven participants (aged 21–50 years, mean = 27.7 years, 4 females) underwent a 90-minutes scan where they were instructed to continuously tap their right-hand fingers for three 10 minutes blocks during the scan. T_2_*-weighted EPIs were acquired during the second finger-tapping block. The total BOLD-fMRI acquisition time was 19 minutes, encompassing the 10-minute task and approximately 4.5 minutes of resting-state data acquired before and after the task block (TR = 2000 ms, TE = 30 ms, voxel size = 3 .125×3.125×3 mm^3^, 37 slices, flip angle = 90°). T_1_-weighted multi-echo MPRAGE were collected (TR = 2530 ms, TE1/TE2/TE3/TE4 = 1.64/3.5/5.36/7.22 ms, TI = 1200 ms, voxel size = 1 mm isotropic, 208 slices, flip angle = 7°).

#### Working Memory Task Dataset

Two independent cohorts of data were included in this study. The first cohort of ten participants (aged 20–47 years, mean = 30.1 years, 7 females) each had a 105 minutes fPET-fMRI scan with 2–4 blocks of resting state sessions and 2–4 blocks of letter-based 2-back working memory task sessions (Chen et al., 2015) with jittered stimulus timings, where T_2_*-weighted EPIs were acquired (TR = 2400 ms, TE = 30 ms, voxel size = 3.125×3.125×3 mm^3^, 44 slices, flip angle = 81°). T_1_-weighted multi-echo MPRAGE were collected (TR = 2530 ms, TE1/TE2/TE3/TE4 = 1.64/3.5/5.36/7.22 ms, TI = 1200 ms, voxel size = 1 mm isotropic, 208 slices, flip angle = 7°). A second cohort of 10 participants (aged 20–38 years, mean = 29 years, 6 females) each had a 28-minute MRI-only functional scan comprised of two 2-minute rest sessions in the beginning and the end, two 8-minute task sessions with 30s-on-30s-off task design, and one 8-minute rest session in between two task sessions, where T_2_*-weighted multi-echo EPIs were acquired (TR = 2000 ms, TE1/TE2/TE3 = 12/27/43 ms, TI = 3, voxel size = 3.4×3.4×3 mm^3^, 26 slices, flip angle = 8°). T_1_-weighted multi-echo MPRAGE were collected (TR = 2510 ms, TE1/TE2/TE3/TE4 = 1.64/3.5/5.36/7.22 ms, TI = 1200 ms, voxel size = 1 mm isotropic, 176 slices, flip angle = 7°). Functional MRI data from two subject cohorts were combined for the computation of graph-theoretical metrics.

#### Sleep Dataset

Fifteen subjects participated in an EEG-PET-MRI scan (mean duration = 87 ± 8 mins), during which they were instructed to close eyes and relax. This dataset is a subset of that reported in our prior study with a different research focus^61^. All 15 subjects included in analysis (aged 23–40 years, mean = 28.9 years, 11 females) were sleep-deprived, instructed to have only 4 hours of sleep the night before the visit to increase sleep pressure and to avoid caffeine on the day of the experiment to broaden the possible range of arousal states. To capture the circadian dip, all experiments were conducted in the afternoon hours. T_2_*-weighted EPIs were acquired (TR = 2400 ms, TE = 30 ms, voxel size = 3 .125×3.125×3 mm^3^, 44 slices, flip angle = 81°) during the scans. Sleep and wake periods were manually identified using sleep index generated from EEG data, and only stable sleep or wake periods lasting longer than 5 minutes were included in the analysis. T_1_-weighted multi-echo MPRAGE were collected (TR = 2530 ms, TE1/TE2/TE3/TE4 = 1.64/3.5/5.36/7.22 ms, TI = 1200 ms, voxel size = 1 mm isotropic, 208 slices, flip angle = 7°).

### Data preprocessing and cortical parcellation

#### fPET-FDG

All fPET images were motion-corrected by registering to the middle temporal frame with *3dvolreg* from AFNI^62^.

#### BOLD-fMRI

All BOLD EPI data went through a standard pipeline of preprocessing: motion correction with *3dvolreg*^62^ and slice timing correction with *3dTshif*t from AFNI^62^, and the working memory and sleep dataset went through distortion correction using an acquisition with opposite phase encoding direction^63^. Nuisance regression was performed for parcellated data and the regressors included were 6 rigid-body motion parameters, low-frequency scan drifts, and the top 5 principal components from white matter and ventricles signals^64^ except for the sleep dataset due to physiological changes during sleep^65^. For visual, motor, and working memory datasets, task-evoked responses were linearly regressed out from task fMRI scans using the convoluted time series of task block design and a canonical hemodynamic response function (HRF) from SPM^66^ and its derivative as regressors.

#### Brain parcellation

Cortical grey-matter regions of interest were delineated based on each subject’s high-resolution anatomical image, using a multi-resolution functional atlas “7 network 300-ROI version”^21^. The Schaefer 300-ROI atlas was projected to each subject’s native surface space for parcellation of preprocessed fMRI data and projected to each subject’s native fPET-FDG space for calculation of ROI-wise CMRglu patterns.

#### CMRglu (PSC)

Task-evoked changes in glucose metabolism were modeled using ramp functions and assessed under a GLM, consistent with prior studies^17,18^. PSC in CMRglu was quantified by dividing the slope of the task portion of the fitted task regressor by the slope of a latter portion of the fitted baseline^22^. For the two constant infusion datasets, we used a linear plus bi-exponential model derived from the tissue compartmental model to characterize baseline^67^; for the two bolus plus constant infusion datasets, we fitted a third-order polynomial to the time series of each ROI to characterize the baseline^18^. The initial 10 minutes of the scan was excluded from analysis to avoid impact of dynamics before tracer equilibrium^18^.

### Meta-analysis records inclusion: resting-state FDG-PET and fMRI studies in disease

Relevant articles from 1983 to 2026 were identified using specific searching strings on PubMed: (1) “(Alzheimer’s disease/Parkinson’s disease/depression/schizophrenia) AND (FDG) AND (PET) AND ((glucose metabolism) OR (cortical metabolism) OR (cerebral metabolism))”; (2) “(fMRI) AND (Alzheimer’s disease/Parkinson’s disease/depression/schizophrenia) AND ((graph theory) OR (degree centrality) OR (nodal degree) OR (nodal centrality) OR (topological metrics))”. A total of 3461 studies were found and evaluated according to a set of exclusion and inclusion criteria.

Studies were included if they 1) were published in a peer-reviewed journal, 2) had adult participants, 3) reported coordinates or provided figures of results from whole brain analysis, 4) were resting-state scans, and 5) were published in English. Studies were excluded if they 1) used other brain imaging modalities, 2) did not use adult human subjects, 3) were review or case studies, or 4) did not include a control group.

After the initial abstract screening, 3,183 of the 3,461 retrieved studies were excluded based on the predefined criteria. The remaining 278 studies underwent full-text screening for eligibility. Additional 188 studies were excluded for the following reasons: did not report regional changes (n = 160), no comparison with healthy controls (n = 26), or duplicate datasets (n = 2). This resulted in 90 studies being included in the meta-analysis. The search procedure is summarized in the PRISMA (Preferred Reporting Items for Systematic Reviews and Meta-Analyses) diagram^68^ (Fig. 1b).

### ROI-based meta-analysis method

We adapted the traditional coordinate-based activation likelihood estimation (ALE) method, which is commonly used in fMRI to identify consistent locations of brain activation reported across studies using similar tasks or states, to access ROI-wise data due to the nature of fMRI-based graph theory studies, and lack of report for coordinates of activation from early PET studies. fMRI-based graph theory studies typically parcellate the brain into bigger ROIs using existing atlases for calculation efficiency, and the AAL atlas with 90 regions^26^ is most commonly used among clinical studies. Hence, we recorded and mapped the results of identified studies on AAL atlas according to their spatial coordinates, if the studies did not adopt AAL atlas as their parcellation scheme. The AAL atlas was projected onto FreeSurfer *fsaverage* surface space, with manual edits applied to refine ROI boundaries.

Among each disorder, we calculated the probability of each ROI having statistically significant alterations in degree or glucose metabolism reported from existing literature. The correlation between spatial probability maps of activation of degree and glucose uptake was assessed with Spearman’s correlation across all cortical ROIs.

### Null models

Null distributions of correlation between complex network metrics and glucose metabolism were constructed through permutation tests with temporal-autocorrelation preserved sham BOLD-fMRI time courses. We generated the sham time courses by randomly reshuffling the phases of the BOLD-fMRI signals in the frequency domain for all subject’s rest and task sessions, calculated the *t*-scores from task/sleep vs. rest complex network metrics difference with the sham data and performed Spearman’s correlation test with respective to true CMRglu PSC data estimated from fPET-FDG. This processed was repeated 5,000 times to derive a null distribution of correlation between complex network metrics and glucose metabolism, and with each other.

### Linking nodal degree to diverse metabolic processes

#### Resting state dataset

A publicly available resting-state simultaneous FDG-fPET/BOLD-fMRI dataset was collected from 27 healthy young subjects in Monash University^69^. Data were acquired on a Siemens 3 Tesla Biograph mMR scanner (Siemens Healthineers, Erlangen, Germany). During the ~ 95-minute scan, participants rested with their eyes open while FDG tracer (mean dose ~ 6.3 mCi) was infused at a constant rate throughout the scan, and T_2_*-weighted EPI images were collected in the meantime (TR = 2450 ms, TE = 30 ms, voxel size = 3 × 3 × 3 mm^3^, 44 slices, flip angle = 8°). Attenuation correction was performed using UTE MRI, and PET images were reconstructed with the OP-OSEM algorithm at a nominal voxel size of 2.09 × 2.09 × 2.03 mm^3^ and a temporal resolution of 16 seconds per frame. Three subjects were excluded due to data quality issues (two with motion artifacts and one with an infusion pump error) as described in our prior study^67^, resulting in a final sample size of 24 subjects (age range 18–23 years, mean = 19.6, 17 females) for analysis. T1-weighted MPRAGE were collected (TR = 1640ms, TE = 2.34 ms, voxel size = 1 × 1 × 1 mm^3^, flip angle = 8°, 176 slices).

#### CMRglu

We quantified the absolute CMRglu by applying the Patlak graphical analysis^70^ using a group-level image-derived arterial input function (IDIF-averaged and smoothed using a cubic smoothing spline with a smoothing tolerance of 0.025×mean signal)^71^ with partial volume correction. The Schaefer 400-ROI atlas, consistent with the parcellation scheme of the shared metabolic pathway data, was registered to each subject’s native fPET-FDG space for calculation of ROI-wise CMRglu patterns.

#### Metabolic pathways

The spatial maps of various metabolic pathways were obtained from a study that used gene sets and whole-brain transcriptomics to map gene expression of metabolic pathways^55^. We assessed the mean gene expression data for 5 major pathways: glycolysis, PPP, TCA, oxidative phosphorylation, and lactate, in our analysis.

To perform correlation analysis with metabolic pathways data, we adopted the same Schaefer 400-ROI atlas for the resting-state dataset to calculate regional degree. Spearman’s correlation test was used to assess the spatial correspondence of resting-state degree and metabolic pathways.

## Acknowledgements

We would like to thank Dr. Mikail Rubinov for helpful discussions of the study findings; Shirley Hsu, Grae Arabasz, Oliver Ramsey, Kyle Droppa, Amy Kendall, David Sosnovik, Marlene Wentworth, and Regan Butterfield for their help with PET-MRI scanning support. We also want to thank Dr. Alain Dagher, Dr. Bratislav Misic, and Mr. Moohebat Pourmajidian for making the cortical distributions of metabolic processes public. This work was supported in part by the NIH (grants K99/R00-NS118120, R01-MH111438, P41-EB030006, and R21-MH135201), by the Harvard Mind Brain Behavior Faculty Research Award, by the Brain & Behavior Research Foundation Young Investigator Grant, by the BrightFocus Foundation Research Grant, by the Rappaport Foundation, and by the MGH/HST Athinoula A. Martinos Center for Biomedical Imaging; and was made possible by the resources provided by NIH Shared Instrumentation grants (S10-RR022976, S10-RR019933, S10-OD010759). Computational resources were generously provided by the Massachusetts Life Sciences Center (https://www.masslifesciences.com/).

