## Supplementary Figures for "Energetic implications of fMRI-based nodal complex network metrics: a complex picture unfolds across diverse brain states"

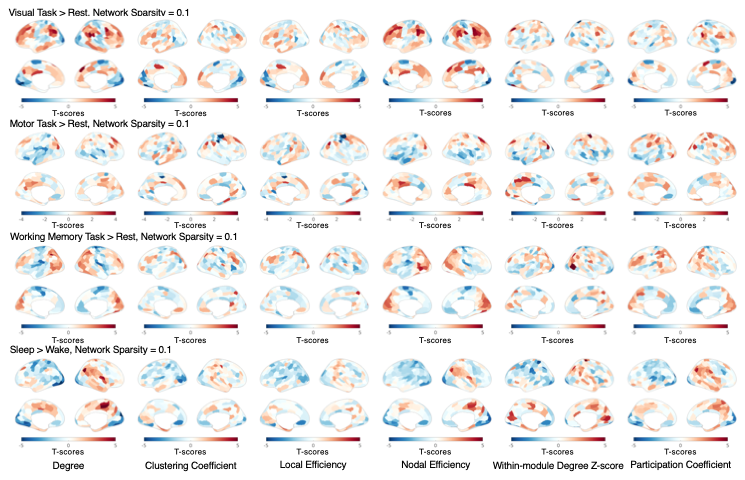

**Supplementary Figure 1. Changes of graph-theoretical metrics across sensory, cognitive, and arousal states.** All functional connectivity matrices were thresholded at network sparsity level = 0.1 (preserving the top 10% strongest functional connections) to calculate the graph-theoretical metrics. Paired sample t-tests were performed to assess the difference between graph-theoretical metrics during task or arousal states and resting state or wakefulness.

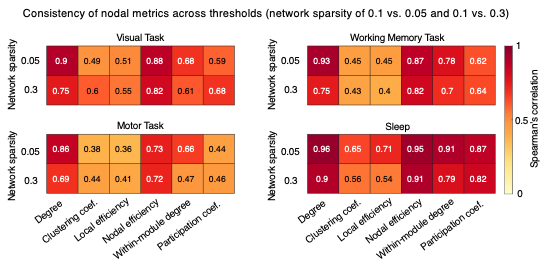

**Supplementary Figure 2. Spatial correlation between nodal graph-theoretical metrics across thresholds.** Spearman’s correlation tests were used to assess the spatial correlation between nodal metrics calculated with network sparsity of 0.1 and nodal metrics calculated with network sparsity of 0.05 and 0.3. Redder color indicates higher correlation.

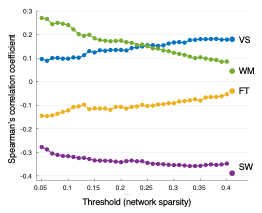

**Supplementary Figure 3. Changes of correlation coefficient between glucose metabolism and nodal degree across functional connectivity matrix threshold in sensory, cognitive, and arousal states.** “VS” = visual task, “FT” = motor task, “WM” = working memory task, “SW” = sleep-wake transitions. All functional connectivity matrices were thresholded at network sparsity level = 0.05-0.4 (preserving the top 5-40% strongest functional connections) to calculate nodal degree. Nodal weighted degree was calculated by adding all functional connectivity strengths. Paired sample t-tests were performed to assess the difference between graph-theoretical metrics during task or arousal states and resting state or wakefulness. Correlation between percent changes in cerebral metabolic rate of glucose and t-scores of changes in nodal degree were assessed with Spearman’s correlation test. Results of weighted degree are highlighted by a bigger marker size.
